# Factors affecting bacterial community dynamics and volatile metabolite profiles of Thai traditional salt fermented fish

**DOI:** 10.1101/2021.02.02.429487

**Authors:** Rachatida Det-udom, Sarn Settachaimongkon, Chuenjit Chancharoonpong, Porrarath Suphamityotin, Atchariya Suriya, Cheunjit Prakitchaiwattana

## Abstract

Bacterial diversity of the Thai traditional salt fermented fish with roasted rice bran or Pla-ra, in Thai, was investigated using classical and molecular approaches. Pla-ra fermentation could be classified into two types, i.e., solid-state fermentation (SSF) and submerged fermentation (SMF). Bacterial population ranged from 10^2^-10^6^ and 10^6^-10^9^ CFU/g in SSF and SMF, respectively. The **rRNA** detection revealed that *Halanaerobium* spp. and *Lentibacillus* spp. were the main genera present in all types and most stages of fermentation. *Tetragenococcus halophillus* were dominant during final stage of fermentation in the samples in which sea salt was used as one of the ingredients while *Bacillus* spp. were found in those that rock salt was used. In contrast, cultural plating demonstrated that *Bacillus* spp. were the dominant genera. *B. amyloliquefaciens* were the main species found in all types of Pla-ra whereas *B. pumilus, B. autrophaeus*, *B.subtili*s and *B. velezensis* were specifically associated with the samples in which rock salt was used. The main volatile metabolites in all Pla-ra samples were butanoic acid and its derivatives. Dimethyl disulfide was observed during earlier stage of fermentation under high salt condition with a long fermentation period. Key factors affected bacterial profiles and volatile compounds of salt fermented fish are type of salt, addition of roasted rice bran, and fermenting conditions.

**Importance:** Protein hydrolysates with high salt fermentation from soy, fish as sauces and pastes are important food condiments commonly found in Asian food cultures. In Thailand, an indigenous semi-paste product derived from salted fish fermentation also called Pla-ra is well recognized and extensively in demands. In-depth information on Pla-ra fermentation ecosystems, in which roasted rice bran and different types of salt are incorporated, are still limited. In this study, we found that *Halanaerobium* spp. was the key autochthonous microbe initiating Pla-ra fermentation. Addition of roasted rice brand and rock salt were associated with the prevalence of *Bacillus* spp. while sea salt was associated with the presence of *Tetragenococcus halophillus*, The risk of pathogenic *Staphylococcus* spp. and *Clostridium* spp. needed to be also concerned. Geographical origin authentication of Pla-ra products could be discriminated based on their distinctive volatile profiles. This research provides novel insights for quality and safety control fermentation together with conservation of its authenticity.

## Introduction

Condiments made by fish fermentation such as fish sauce, shrimp paste, and fish paste are important food products commonly found in Asian, particularly Southeast Asian countries. In Thailand, products like fish sauce, fish paste, and semi paste (Pla-ra) are commercially manufactured. Marine fish have been used for sauce production while freshwater fish have been used in the other fermented products, mainly Pla-ra.

Recently, demand of Pla-ra in both domestic and export markets have been increased since Thai fusion dishes from exotic ingredients are widely created. Pla-ra is used as condiment, thickening sauce, dipping paste and snack and becomes popular.

After fermentation Pla-ra contains both fine meat texture and thickening meat-derived liquid providing Kokumi taste (1) with fishy and volatile metabolite aroma. Pla-ra manufacturing is significantly different from fish sauce. It is (i) traditionally produced from variety of natural freshwater fish, locally and seasonally harvested in the local area; (ii) preserved, preferably with partially purified rock salt; (iii) produced with addition of roasted rice bran; and (iv) fermented for 8-24 months depending on manufacturing process. Thus, Pla-ra characteristics from each production area are unique.

Microbes associated with salt fermented fish, particularly fish sauce, are well investigated and their role in fish protein hydrolyzation including metabolic activity are well known. With indigenous enzymes from fish, microorganisms during an initial period of fermentation such as *Halanaerobium*, *Bacillus*, or *Staphylococcus*, utilize raw materials and change nutrient molecules to primary metabolites of amino acids, glucose and fatty acids via their proteolytic and lipolytic activity (2). These substances then support the growth of the other microbes such as *Halomonas*, *Tetragenococcus*, and *Trichococcus* in subsequent fermentation stages, and generate various sensory compounds (3, 4).

These microbial community are also similar to the microbial profile in another fermented fishery product such as Korean salted and fermented seafood called Jeotgal (5, 6), salt-fermented shrimp paste (1) and anchovy sauce Budo (7). Even though the comparable trends of bacterial profiles were proposed, many environmental factors including raw materials, formulation, equipment and production process were highly affected microbial community dynamics during fermentation (2).

Similarly, the Pla-ra fermentation may include the main steps of fish protein fermentation under high salt concentration. However, it is a challenge to investigate whether freshwater fish, rock salt and particularly, roasted rice bran affects microbial communities and fermentation activities under high salt condition. The information obtained could provide new insights during Pla-ra fermentation regarding the influence of types of fish, key ingredients and preparation methods on microbial population dynamics and volatile metabolite formation.

Upon this, technologically relevant characteristics as well as molecular authenticity of the product could be established. Since the domestic and international trade demands of Pla-ra extensively increase, investigation of these parameters provides a great opportunity to further develop production technologies for better quality and safety control. The originality of this indigenous product could also be well-conserved. This research systematically investigated influence of key factors including production area and manufacturing conditions on culturable and non-culturable bacterial community. The physiochemical properties and volatile metabolite profiles associated with each bacterial ecosystem of Pla-ra during fermentation were also studied.

## Results and discussion

### Production area and manufacturing process

Data were collected in selected regions regarding manufacturing process and sources of raw materials of Pla-ra that might affect its characteristics. The manufacturing process and raw materials of Pla-ra were predominantly influenced by production area and local culinary culture (Table S1).

Tilapia (*Oreochromis niloticus*) caught from Mekong river was the main large-size fish used as a raw material in the production of Pla-ra in A1 (Udon Thani) and A2 (Nongkai) provinces (Figure 1). However, a variety of small-size fishes such as catfish (*Mystus cavasius*) and Henicorhynchus (*Henicorhynchus siamensis*) were occasionally used for the production of homemade products. Pla-ra made in A1 and A2 provinces was fermented in solid state for > 6-12 months with addition of approximately 10% of roasted rice bran and 20-25% of rock salt. Rock salt used in this area was produced from Bandung district (A1.3) in A1 province.

**Figure 1.**
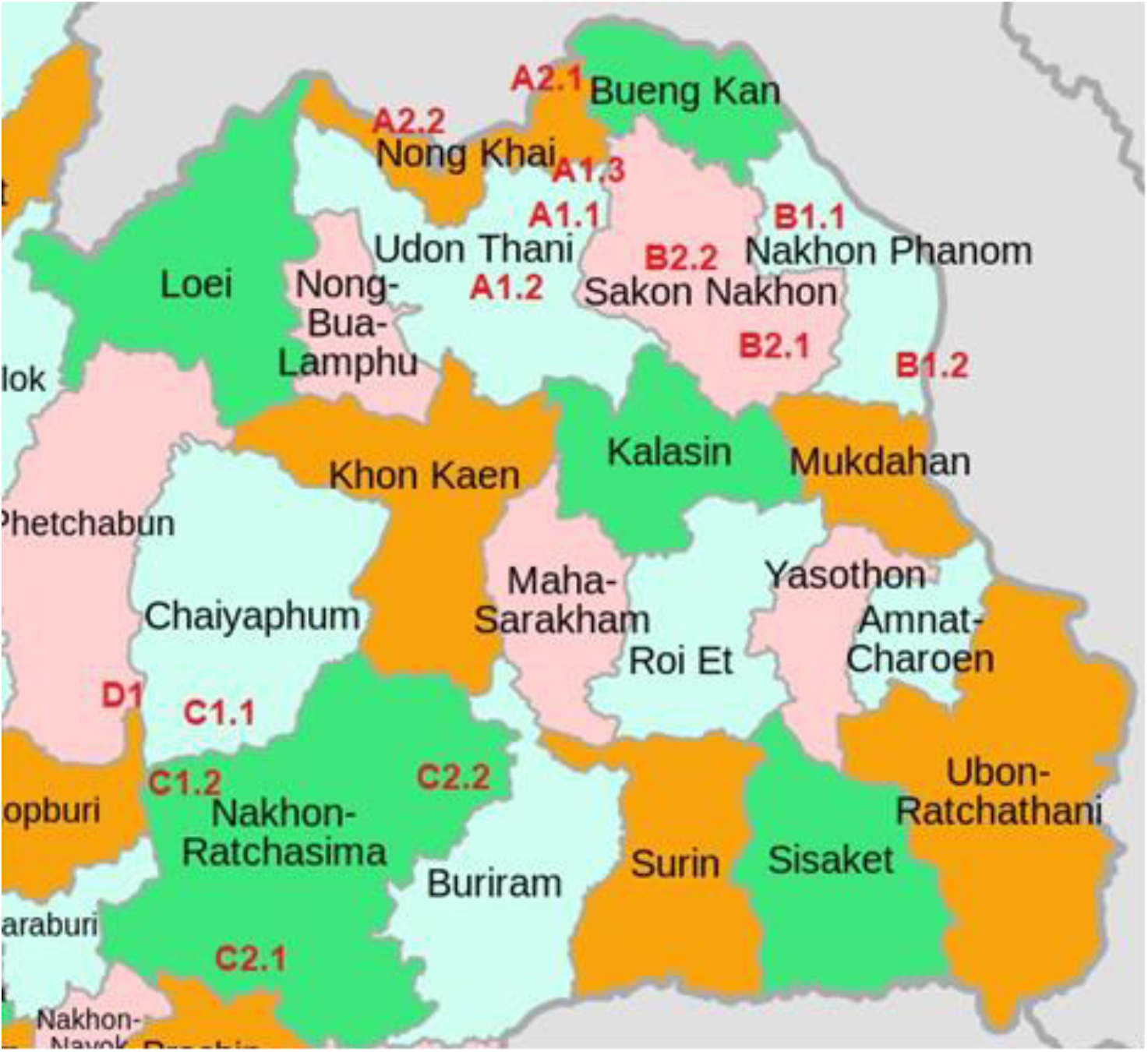
Map of manufacturing sites and source of raw materials used in this study

In B1 (Sakon Nakhon) and B2 (Nakhon Phanom) provinces, a great variety of small-size fishes, including *Mystus cavasius* and *Barbus gonionotus* (Thai carp) etc., caught from local reservoirs, were the main fish used in the production of Pla-ra. A long duration of solid-state fermentation for over 8-24 months was carried out with low amount of rice bran and extremely high amount (up to 25-35%) of rock salt from different origins depended on their availability. Colour of Pla-ra produced in this area were typically darker compared with those from the other provinces

Unlike the two previous areas, cured fishes from reservoir of Pasak Jolasid Dam (D1) located in central Thailand, were transported to C1 (Nakhon Ratchasima) and C2 (Chaiyaphum) provinces for production. Cyprinidae (*Henicorhynchus siamensis*) partially mixed with Anabantidae (*Anabas testudineus*) and Gouramis (*Trichopodus* spp.) were considered as the main ingredient. The fishes were usually cured under >40% salt for 2-4 weeks before selling to Pla-ra manufacturers. It should be highlighted that an addition of water at a ratio of 1:1 was required for reducing their saltiness prior to Pla-ra fermentation when cured fishes were used. Accordingly, a submerged fermentation system with addition of >30% roasted rice bran and 17-22% sea salt, sometimes in combination of rock salt, were mainly used for a duration of usually less than 6 months.

### Bacterial profiling during Pla-ra fermentation using classical and molecular approaches

Bacterial population dynamics during Pla-ra fermentation were characterized using cultural plating technique in combination with **rRNA** transcriptional analysis (Table 2). Overall results from the **rRNA** analysis revealed that *Halanaerobium* spp. and *Lentibacillus* spp. were the two major bacterial population dominating in Pla-ra samples from all production regions at all stages of fermentation. Based on the cDNA intensity, the *Halanaerobium* spp. showed a higher **rRNA** level (high-intensity cDNA band) compared to *Lentibacillus* spp. during early fermentation period (1-3 months). **rRNA** of the two genera were detected at similar level during the mid-period of fermentation (4-9 months) and almost disappeared from the fermentation ecosystem at the end of fermentation (>9 months).

During different stages of fermentation, *Lactobacillus acidipiscis* and *Staphylococcus* spp. were detected specifically during early fermentation when multi-species of fish were used as raw material in the area of A1 and A2. Also, *Bacillus* spp. was detected in the samples from A2-2 which were fermented over a year, while it was not found in those with shorter fermentation period.

Even though *Halanaerobium* spp. and *Lentibacillus* spp. were also the two major genera in the samples collected from regions A and B, it was found that *Lentibacillus* spp. was replaced by *Tetragenococcus halophillus* particularly in samples fermented over a year and various species of fish were used as raw material in combination with sea salt in region B. Nevertheless, the **rRNA** of *Bacillus* spp. and *Staphylococcus* spp. genes were not detected in samples collected from B.

The effect of salt used in the process was further investigated. It was found that **rRNA** of *Lactobacillus acidipiscis* and *Lactobacillus* spp. were also detected in samples fermented over a year with addition of rock salts produced within region A.

Consistent with the results from regions A and B, *Halanaerobium* spp. and *Lentibacillus* spp. were also the two major genera detected in Pla-ra samples collected from C1 and C2. RNA level of *Halanaerobium* spp. was higher compared to *Lentibacillus* spp. throughout the entire course of fermentation. It should be mentioned that Pla-ra production in region C is generally performed by submerged fermentation (SMF) with shorter incubation period (<6 months) compared to solid-state fermentation (SSF). This short duration could be considered as early stage of fermentation in the other regions where *Halanaerobium* spp. and *Lentibacillus* spp. were both detected as major species dominating the microbial ecosystem.

Besides Pla-ra from the major production regions of Thailand, representative samples from Ubon Ratchatani and Mukdahan, Thailand as well as Vientiane, Lao People’s Democratic Republic were also investigated. Bacterial **rRNA** results confirmed that *Halanaerobium* spp. and *Lentibacillus* spp. were the predominant genera present in Pla-ra fermentation ecosystem (data not shown). The **rRNA** of *Clostridium* spp. and *Staphylococcus* spp. were clearly observed, especially in the samples containing more than 18% salt. Further investigation is required for food safety management since both are pathogens which shall be controlled to minimize public health risk.

### Cultural plating

The number of bacteria ranged from 103-107 CFU/g with only small diversity of colony morphology observed (Table 1). The number of bacteria in Pla-ra samples collected in the regions A and B using SSF (103-106 CFU/g) were lower than those detected in samples collected in region C (106-107 CFU/g) where SMF was employed (Table 2). The number of bacteria tended to decrease towards the end of SSF.

**Table 1.** Characteristics of Pla-ra samples collected from various regions of Thailand

| Areas | pH /<br>TTA /<br>Salt (%) | color<br>(L / a / b) | Proximate analysis*<br>(% by weight)<br>(MS / ash / fat / fiber /<br>protein) | Microbial population**<br>(approx.Log CFU/g) |  |  |
| --- | --- | --- | --- | --- | --- | --- |
|  |  |  |  | TVC | Y&M | SA |
| A1 / A2 | 5.44 ± 0.44 / | 45.44 ± 6.58 / | 64.62 ± 3.03 / 7.54 ± 2.14 / |  |  |  |
|  | 1.49 ± 0.31 / | 4.34 ± 2.07 / | 8.85 ± 2.02 / 4.98 ± 1.24 / | 4.00-5.33 | 3.40-4.70 | 0.13 |
|  | 24.53 ± 1.35 | 12.07 ± 4.37 | 11.06 ± 1.22 |  |  |  |
| B1/ B2 | 5.4 ± 0.98 / | 47.79 ± 6.43 / | 61.32 ± 6.40 / 12.77 ± 3.31 / |  |  |  |
|  | 1.54 ± 0.34 / | 5.66 ± 3.29 / | 9.23 ± 0.75 / 3.79 ± 1.21 / | 3.86-6.30 | 1.00-2.30 | 0.22 |
|  | 32.43 ± 2.25 | 10.58 ± 4.17 | 9.76 ± 3.41 |  |  |  |
| C1 / C2 | 5.89 ± 1.11 / | 49.39 ± 5.76 / | 64.57 ± 1.52 / 6.62 ± 0.11 / |  |  |  |
|  | 0.67 ± 0.29 / | 7.37 ± 2.63 / | 5.63 ± 1.04 / 12.1 ± 1.06 / | 6.00-6.74 | 1.00-3.88 | 0.13 |
|  | 21.67 ± 1.73 | 15.87 ± 4.47 | 12.1 ± 1.26 |  |  |  |

\* MS = moisture content, TTA = Tritatable acidity (based lactic acid) H = Hardness, C = Cohesiveness
\*\* TVC = Total viable count, Y&M = Yeast &Mold count, SA = *Staphylococcus aureus*

**Table 2.**
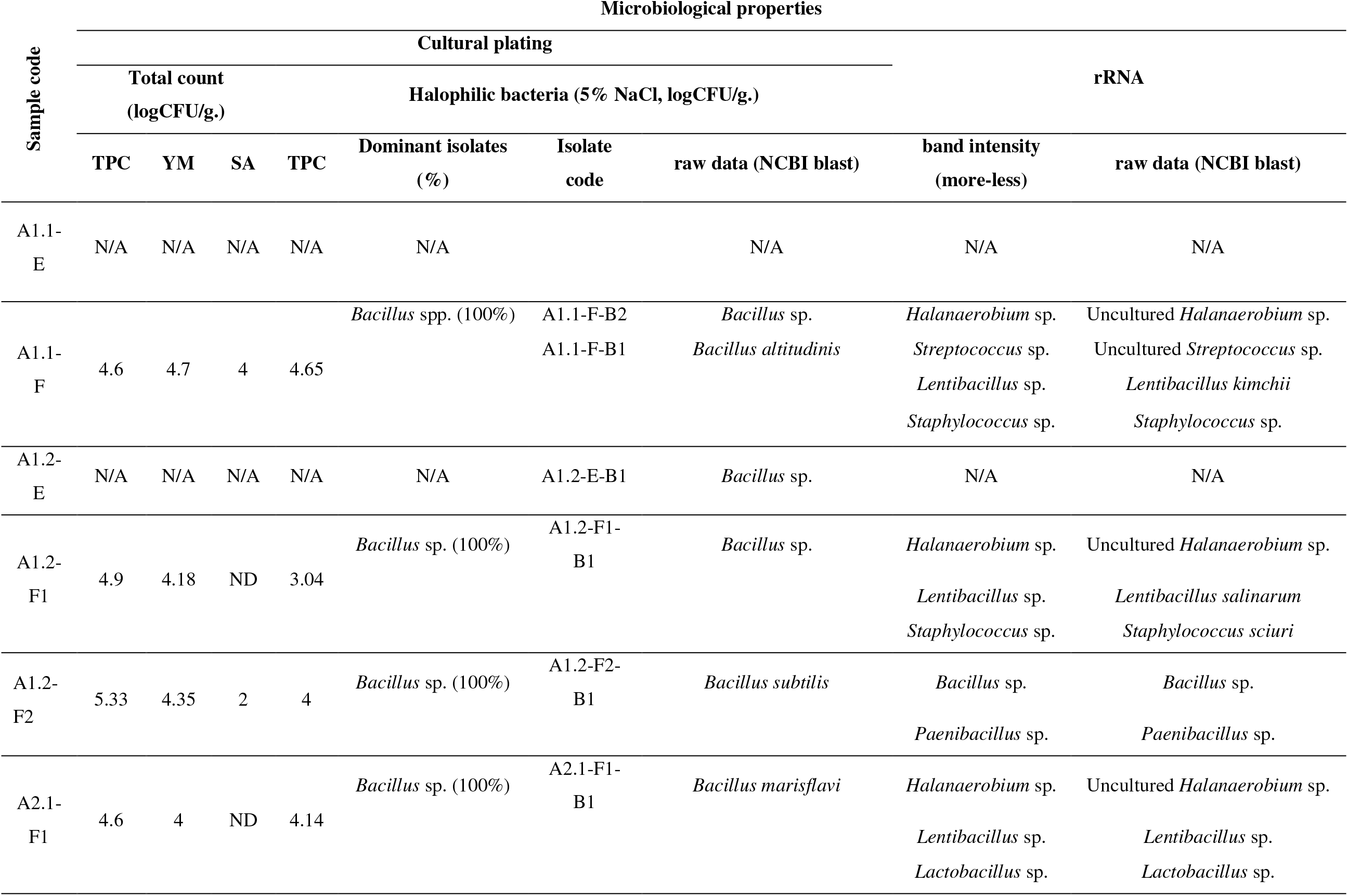
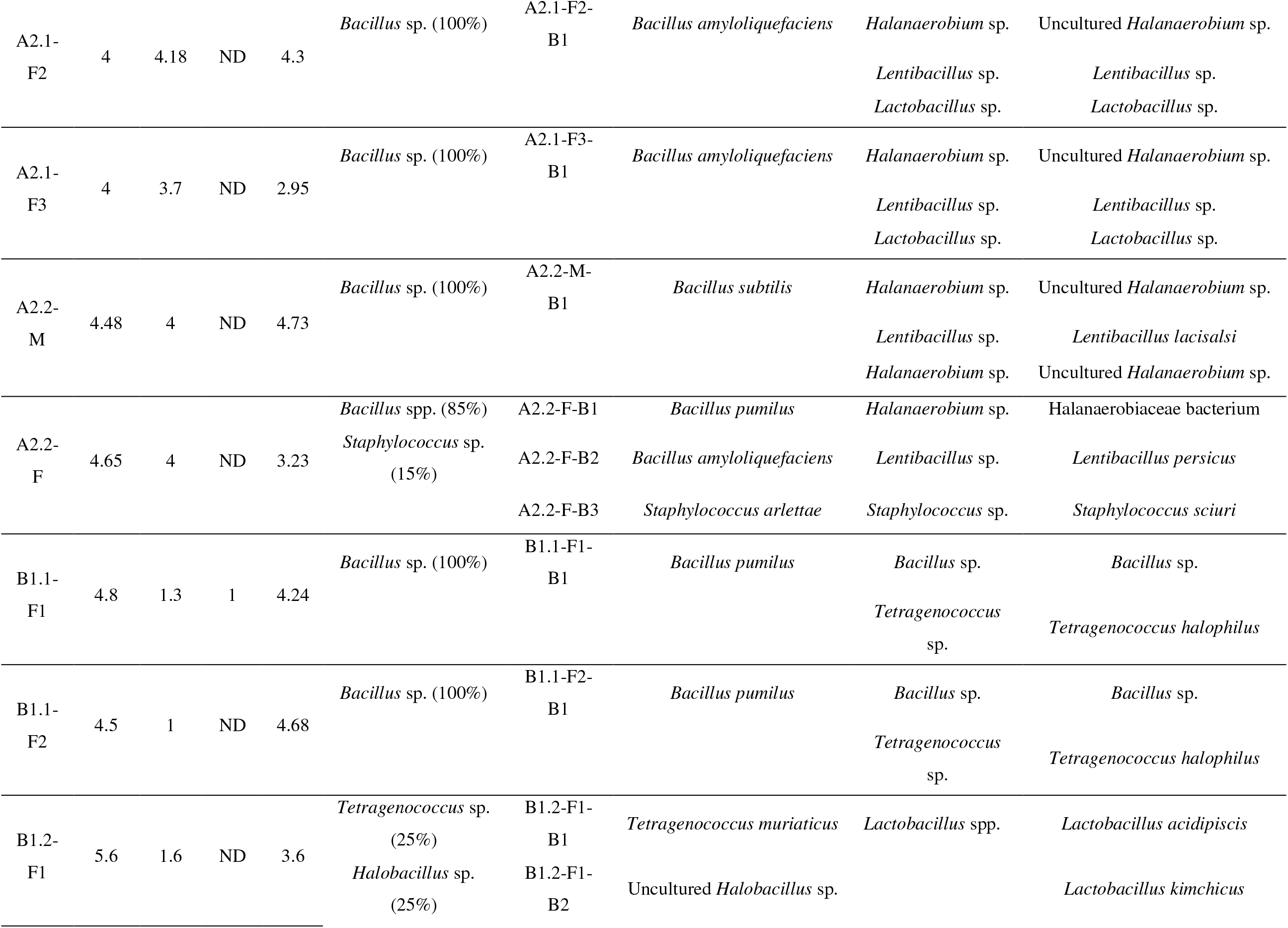
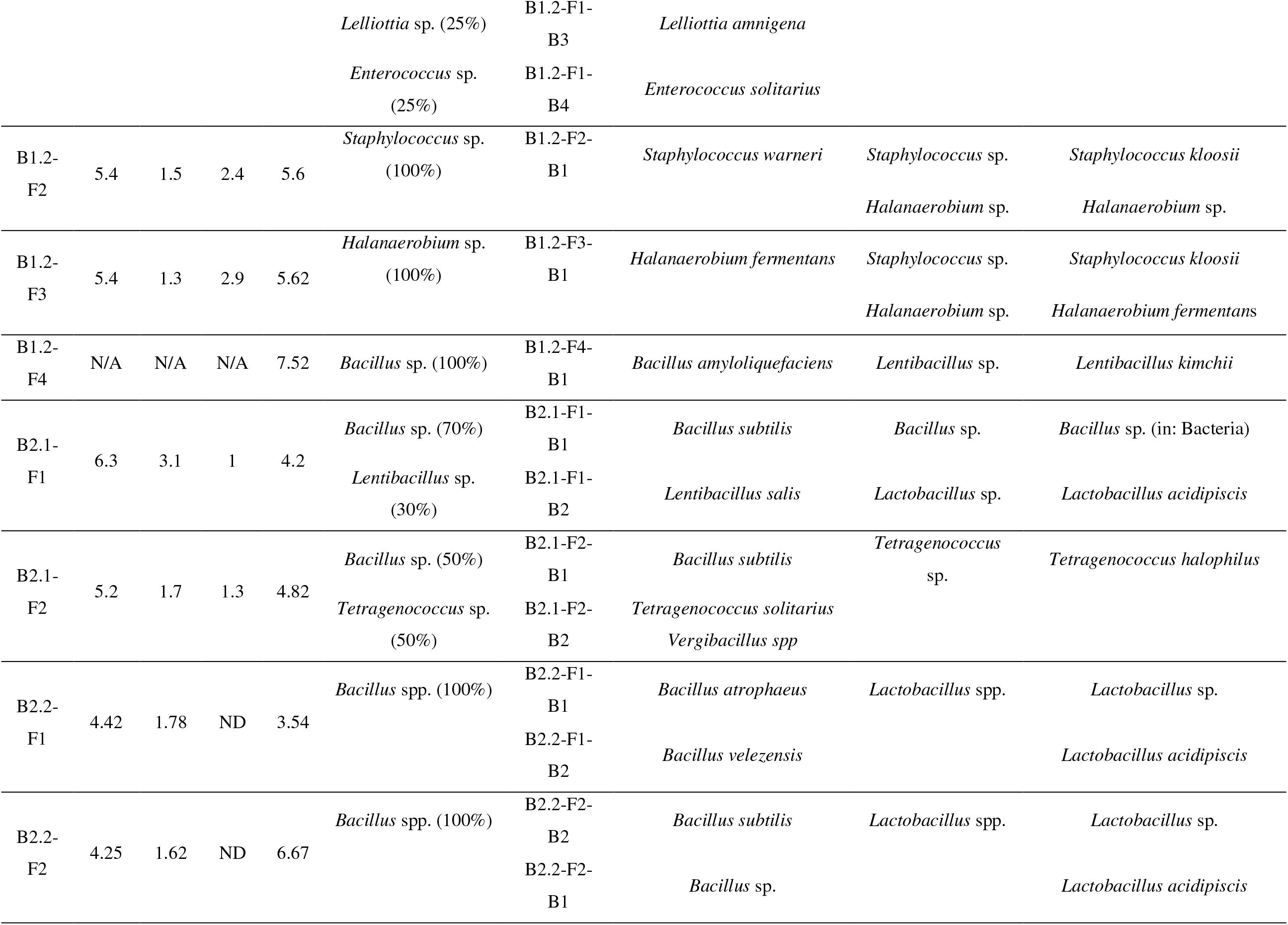
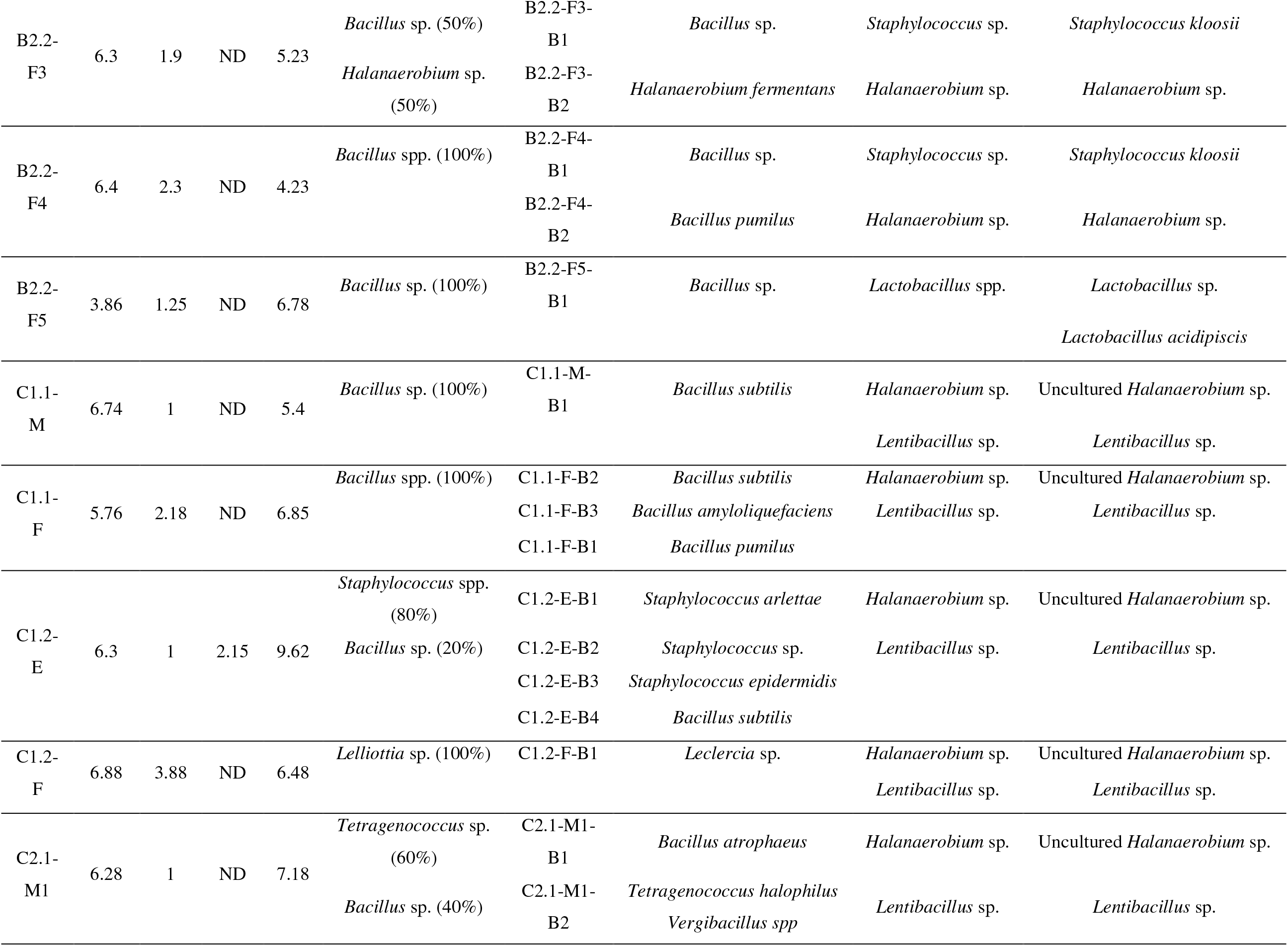
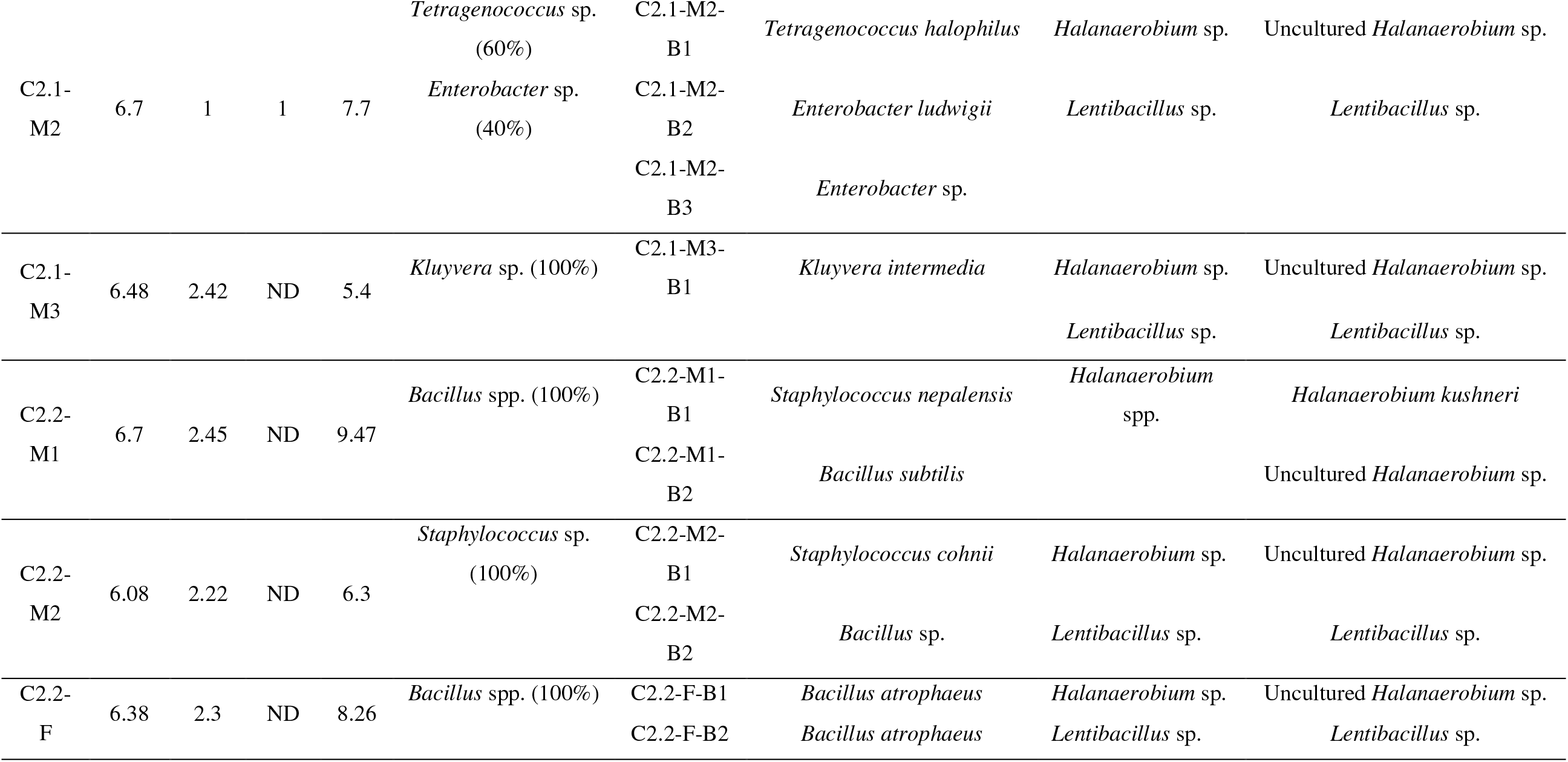
Bacterial population profiles of Pla-ra samples

In contrast, the bacterial community identified by classical method differed from that of the molecular approach. The result revealed that *Bacillus* spp. was the main isolate found in most Pla-ra samples collected from all regions. *Staphylococcus* spp. was also found in some samples.

Interestingly, *T. halophilus* was the main isolate in the samples collected in region C which were made from cured fish with added sea salt while *Bacillus* spp. was the main isolate in those made from fish caught within the regions of A and B.

In samples fermented over six months with addition of sea salts, more diversity of microbial community was found in the samples from region B. Besides *Bacillus* spp., *Tetragenococcus halophillus*, *T. muriaticus*, *Vergibacillus* spp., *Lelliottia* spp., *Halobacillus* spp., *Oceanobacillus* spp. and *Lentibacillus* spp. were identified.

The domination of *Bacillus* spp., a genus with endospore forming capacity, could be associated with the application of roasted rice bran which was one of the ingredients in the manufacturing process. The rice bran could be a good source of bacillus spores which can later germinate and play important roles due to their amylolytic, proteolytic and metabolic activities during SSF.

However, the expression of *Bacillus* genes was not detected by **rRNA** analysis since very small amount (1-10%) of roasted rice bran, the suspected source of *Bacillus* spp., was added compared to the source of *Halanaerobium* spp. and *Lentibacillus* spp which was the GI tract of the fish and salt (14). The initial population of *Bacillus* spp. might be therefore lower than the latter two bacteria. Thus, the cDNA of *Bacillus* might not be sufficiently primed and amplified if it was significantly lower than the first two prevalent populations (15). However, fresh fish naturally contains nitrate, so it could support *Bacillus* to grow anaerobically by respiration with nitrate (16), allowing these bacteria to ferment as observed by cultural plating.

The prevalence of *Halanaerobium* spp. throughout the entire fermentation period was in agreement with the work of Kobayashi et al. (17) who found most strictly anaerobic bacteria such as *Clostridium* spp., *Halanaerobium* spp. and a variety of halophilic lactic acid bacteria were associated with salted fish fermentation systems. *Halanaerobium* spp. is a halotolerant species which can grow in an extremely high salt environment (>20%) (3). It has been documented that *Halanaerobium* spp., a strict anaerobe, are responsible for the conversion of thiosulfate to sulfide accounting for the dark colour and unique flavour of fermented fish (18).

The results also demonstrated that *Lentibacillus* spp. was predominantly observed in salt fermented freshwater fish under limited oxygen concentration. *Lentibacillus* spp. has also been reported as an extreme halophile with endospore forming capacity, bacterium associated with salted seafood fermentation systems such as anchovy, shrimp paste and fish sauce (19, 20).

It should also be mentioned that the prevalence of *Tetragenococcus halophillus* seemed to be associated with the application of sea salts in the recipe of region B and C.As a result, the mineral contents of rock and sea salts were additionally analysed. It was found that the concentration of potassium (K) in sea salt was approximately 5-40 times higher than those present in rock salt (data not shown). It has been documented that K is essentially required and widely applied as supplement in certain selective media, in order to stimulate optimal growth of *Tetragenococcus* spp. (21). This finding is relevant and requires further investigation since supplementation of K-rich salts might be used to promote the development of *Tetragenococcus* spp. in Pla-ra fermentation. Their active growth and metabolic activity have been associated with the formation of various metabolites responsible for desirable flavour of Pla-ra as well as fermented shrimp paste, miso, fish sauce and soy sauce products (11, 17, 22).

### Phylogenetic trees

To find the relationship among isolates that might reflect key characteristics in Pla-ra fermentation, phylogenetic trees were constructed from 16s rRNA bacterial isolate (Figure S1 A to C). The first largest group was *Bacillus*. Even though *Bacillus* is generally reported as a dominant autochthonous in salt fermented fish and its related products (3, 6, 7), a vast genetically difference among their 16s rRNA sequences were clearly remarked in our study with a few preserved strains from A1 and A2, and C1 and C2. The second group was *Staphylococci*. The tree clearly depicted a significant heterogeneity in their sequences even a common ancestor was shared. In the third group of multi species, a certain divergence of genealogical relationship was depicted, with their root of *Enterobacter*.

Focusing on bacterial expression of the dominants, their phylogenetic variations between production sites were found. Metabolically active *Halanaerobium* spp. and *Lentibacillus* spp. strains detected from Chaiyaphum and Nakorn Ratchasima samples which sea salt was applied generally shared the same internal node apart from others. This reflected a genetically diversity between autochthonous bacteria playing roles in Pla-ra fermentation from different regions and/or formulations.

### Physicochemical property of Pla-ra

By sensory evaluation, desirable characteristics of Pla-ra are described by the characteristics of (i) its liquid portion which should be dark and viscous, and (ii) its solid portion which should be red and paste-like, with high intensity of fish sauce smell (salty, fishy, stinky) and umami taste. All samples in this study shared these distinct quality aspects.

Determination of colour showed no significant difference in *L\**, *a\** and *b\** values among samples (Table 1). It could be noted that Pla-ra’s colour became darker during the course of fermentation (8-24 months). In case of C1 and C2 provinces, roasted rice bran was added to darken the products due to their short fermentation period.

Regarding acidity, the pH values of Pla-ra produced in C1 and C2 provinces (5.89±1.11) were significantly higher compared to those observed in the samples collected from A1 and A2 (5.44±0.44) and B1 and B2 (5.40±0.98). The total titratable acidity (TTA) of samples also corresponded well with their pH values. The acidity seemed to be lower with longer period of fermentation. This differences in acidity among samples seemed to be associated with the duration of fermentation and type of salt used. According to our results, the presence of lactic acid bacteria (i.e. *T. halophillus*) which caused higher level of acid were significantly associated with sea salt. Also, the prevalence of lactic acid bacteria was mostly at the final stage of fermentation. Thus, the samples fermented with sea salt and/or at later stage of fermentation was found to contain higher acidity.

Based on information obtained in this study, type and composition of fish were not likely the key factors affecting the microbial community of Pla-ra. On the other hand, type of salt and fermenting conditions seemed to have an influence on microbial community which resulted in key microbes playing an important role in proteolytic and metabolic functions during Pla-ra fermentation.

### Key factors influencing volatile metabolite profiles during Pla-ra fermentation

The volatile metabolite profiles of Pla-ra samples were characterized and compared using a non-targeted GC/MS-based metabolomics combined with multivariate analysis (Figure 2). Results demonstrated that organic acids, especially butanoic acid (rancid-buttery flavour) and a series of butanoate esters (fruity, buttery, cheesy, greeny flavour), aldehydes, as well as several sulphur containing compounds, especially dimethyl disulfide (sulfurous, vegetable-like flavour) and dimethyl trisulfide (sulfurous, meaty, greeny, onion-like flavour) were the most abundant metabolites present in Pla-ra samples. This finding is consistent with the volatile profiles previously determined in another fermented fish products (4, 23). It is well documented that indigenous enzymatic degradation of fish flesh and microbial activities, especially hydrolysis and metabolism of proteins and lipids, play an important role in flavour development during fish fermentation (24).

**Figure 2.**
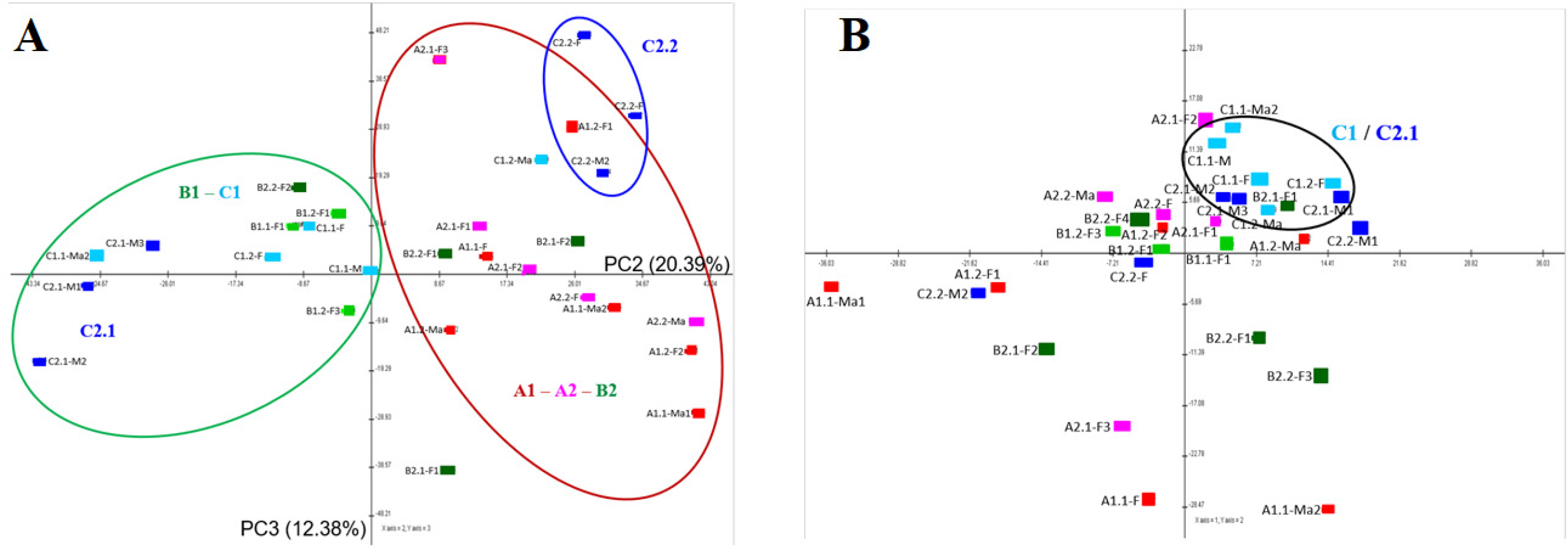
PCA analysis of potential biomarkers in Pla-ra. A = volatile metabolites; B = amino acid profiles

Based on the RNA results, *Halanaerobium* spp. seemed to dominate microbial population especially during the early fermentation stage. It has been reported that *Halanaerobium praevalen*s and *Halanaerobium alcaliphilum*, were able to produce CO_2_ and a number of metabolites, i.e. lactate, acetate, propionate, butyrate and various sulphur compounds, that predominantly contribute to the specific malodorous characteristics and brownish colour of the fermented product (25). This finding was in agreement with the work of Jung et al. (26) who stated that the presence of *Halanaerobium* could be a potential indicator for off-flavour development in fermented shrimp and seafood.

In addition, the presence of *Staphylococcus*, *Virgibacillus* and *Tetragenococcus* are usually observed during the production of fermented seafood (24). These bacteria have an important role in flavour characteristic of product attributed to their lipolytic and proteolytic activities (24, 27). A positive relationship between the presence of *Virgibacillus* and *Tetragenococcus* and the generation of glutamyl peptides responsible for taste enhancers of Pla-ra has been acknowledged (1). It was also reported that an increase of *Staphylococcus* was accompanied by the development of esters in relation with their high catalytic activity (4, 28).

*Bacillus* was observed as the main culturable bacteria isolates in Pla-ra. The presence of these halotolerant bacilli is due to their ability to form endospores to survive under prevailing conditions (29). The halotolerant bacilli have strong influence on metabolism of proteins due to their proteolytic activity (24).

Besides effect from diverse microbes, it should be noted that the flavour metabolites such as amino acids, oligopeptides, organic acids, amines and esters could be varied due to the fish used as raw material, concentration of ingredients and dynamics during different fermentation stages (4). Our results demonstrated that the type and proportion of rice bran and concentration of salt significantly influenced the volatile metabolite fingerprints of Pla-ra samples. It has been reported that different enzymes were activated and the type and activity of microbes changed at different salt levels, resulting in different end products (30).

Unsupervised pattern recognition was performed using Principal Component Analysis (PCA) in order to determine the overall biomolecular characteristics of Pla-ra in association with production area and manufacturing process (Figure 2A). Results demonstrated that samples could be predominantly classified into four groups based on their volatile metabolite profiles, i.e. (i) Pla-ra collected from A1, A2 and B2 provinces, (ii) Pla-ra collected from B1 and C2 provinces, (iii) and (iv) Pla-ra collected from different districts in C1 province. It should be noted that the two SMF samples between the two districts from province C are completely distinguished from each other. Loadings of PC2 indicated potential biomarker metabolites accountable for the discrimination (Figure 2A).

The concentration of 3-methyl-1-butanol, 3-methyl butanoate, 1-octen-3-ol, acetone, benzaldehyde, hexanal and 3,7-dimethyl-1-octanol were present in higher relative abundances in samples collected from A1, A2, B2 and a part of C1 areas compared to those collected from B1 and C2 provinces. It was found that rock salt was used as a major ingredient in the manufacturing process of these samples. In contrast, the concentration of butanoic acid, ethanol, ethyl-butanoate, 3-methyl-butyl-butanoate, phenol, propyl-butanoate, dimethyl disulfide, 1-methyl-ethyl-butanoate, 4-methyl-ethyl-butanoate, 1-butanol and butyl-butanoate in samples collected from B1, C2 and a part of C1 provinces were present at higher relative abundances compared to those detected in A1, A2 and B2 groups. Sea salt was primary used as a major ingredient during Pla-ra production in these regions. This finding demonstrated a possible molecular-based geographical authentication of Pla-ra products based on their volatile metabolite profiling.

Regarding non-volatile metabolites, amino acid profiles of samples were also analysed to investigate the relationship among bacterial composition, proteolytic activity, amino acid metabolism as well as volatile metabolite profiles. A PCA score plot was constructed based on the relative concentrations of 20 amino acids determined by HPLC technique (Figure 2B). Unlike volatile metabolite profiles, different pattern recognition of amino acid profiles could not be observed among Pla-ra samples. As a result, the influence of neither geographical origin nor manufacturing process could be predicted.

In summary, this study demonstrated that the bacterial dynamics of Thai traditional salt fermented fish with roasted rice bran were mainly subjected to the manufacturing process, particularly salt type and fermentation period. Cultural plating demonstrated diverse *Bacillus* at all stages of fermentation while **rRNA** revealed that *Halanaerobium*, *Lentibacillus. Tetragenococcus halophillus* were dominant during final stage of fermentation when sea salt was used and *Bacillus* spp. were only found in rock salt fermentation.

## Materials and Methods

### Survey and Sample collection

Salt fermented fish with roasted rice bran called Pla-ra manufactured in 12 districts from 6 Provinces of the north eastern Thailand, as shown in Figure 1, was investigated. Pla-ra samples were collected from different manufacturers, processes and fermentation stages. Different fermentation stages were early (E), middle (M), final (F) and matured (Ma) (Table S1, supplementary data), depending on manufacturing process of each area. All samples were collected and transported to the laboratory in sterile closed containers. Physical and chemical properties such as texture, colour, proximate analysis, pH, total titratable acidity (TTA) and salt concentration as well as microbial community and metabolomic/volatile compounds pattern were determined. The bacterial communities were investigated and isolated with both cultural dependent and gene expression (**rRNA)** analysis.

### Bacterial community and identity assays

For the cultural independent method, reverse transcriptase-PCR-denaturing gradient gel electrophoresis (PCR-DGGE) was used following Chhetri et al. (8) with some modification. The first universal bacterial primer set was 27F and 1492R (9) and the second was 357F with GC clamp attached and 517R (10). Amplification was done in a standard reaction mixture (Taq DNA Polymerase, Vivantis, Malaysia) following manufacturer’s instruction in a DNA thermal cycler (BioRad T100TM Singapore). DGGE analysis was performed following electrophoresis technique, DNA bands was then extracted, reamplified and subjected to DNA sequencing analysis.

The cultural dependent method was conducted by spreading serial dilutions of liquid samples onto Nutrient agar (Himedia, India) supplemented with 5% NaCl and incubated at 37 °C for 48 h. The colonies were counted and selected by the Harrison’s disc method and colony morphologies. The salt tolerance and protease properties of isolated colonies were screened according to Tanasupawat et al. (11) and Sánchez-Porro et al. (12), respectively. The isolates were identified by DNA sequencing analysis, targeting conserved regions of the 16S rRNA V3. PCR was performed using primer set of 338F/518R (8). The sequencing data was analysed with nucleotide BLAST program of NCBI

Phylogenetic and molecular evolutionary analyses were conducted. Based on alignments of sequencing results of isolates with Basic Local Alignment Search Tool Nucleotide (BLASTN) (www.ncbi.nlm.nih.gov), cultures showing the homologous 16S rRNA gene sequence were selected for phylogenetic analysis. Phylogenetic relationships were examined using MEGA X. These phylogenetic tree reconstructions were carried out using the maximum-likelihood method and the neighbour-joining algorithms with 1000 randomly selected bootstrap replications. The bootstrap consensus trees were constructed based on a topology of the most frequently appearing branch groupings.

### Analysis of volatile metabolites using headspace SPME-GC/MS

About 3 g of Pla-ra was weighed into 20 ml of headspace vial and capped. Sample was pre-heated at 40 °C for 10 min, and a SPME fibre (50/30μm DVB/CAR/PDMS, SUPELCO, PA) was then used to extract volatile compounds at 40 °C for 30 min. The fibre was desorbed in GC injector port at 250 °C for 5 min. Separation of the desorbed volatiles was achieved by gas chromatography–mass spectrometry (Agilent 7890A GC-7000 Mass Triple Quad) equipped with a capillary column (DB-WAX, 60 m × 0.25 mm × 0.25 μm, J&W Scientific, Folsom, CA) and a quadrupole mass detector. The injector was operated at split mode with a split ratio of 5:1. Helium gas was used as the carrier gas with a constant flow rate of 0.8 mL/min. The GC oven temperature was started at 32 °C for 10 min, increased to 40 °C at 3 °C/min and hold for 15 min, then increased to 160 °C at 3 °C/min, then increased to 230 °C at 4 °C/min and hold for 5 min. The mass spectrometer was used in the electron ionization mode with the ion source temperature set at 230 °C, and ionization energy set at 70 eV. The scan mode was used and the scan range was 25 to 400 m/z. The Agilent Mass Hunter Qualitative Analysis B.04.00 software was used for data analysis. Identification of volatile compounds was performed by comparing mass spectra with NIST mass spectral libraries (National Institute of Standards, 2011 version). The content of volatile compound was calculated from peak area.

### Amino acid profile analysis

Amino acid (AA) profile analysis of samples was performed using high-performance liquid chromatography (HPLC). Samples were filtered through a 0.45 μm nylon syringe filter. Then, a derivatization with o-phthalaldehyde was performed. Free AA profile was analysed using an HPLC equipped with a UV-VIS detector (Agilent 1100, USA) at 338 nm. Free AAs were separated on a C18 (ZORBAX Eclipse-AAA, 4.6 × 150 mm, 5 μm) with a guard column, at 40°C. The gradient elution system consisted of a mixture of 40 mM sodium phosphate dibasic, pH 7.8, and a mixture of acetonitrile : methanol : water (45:45:10 v/v/v). The gradient program was set as follows: 0–1.9 min, 0%B; 18.1 min, 57%B; 18.6 min, 100%B; 22.3 min, 100%B; and 23.2 min, 0%B and then held for 2.8 min to re-equilibrate for initial condition before the next injection. The total running time was 26 min with the flow rate of 2.0 mL/min. AAs in Pla-ra samples were identified by comparing the retention time with AA standard. Quantity of AA (mg/100 mL) was determined based on the external standard method using calibration curves fitted by linear regression analysis.

### Statistical analysis

Analysis of variance and multiple comparisons by Tukey’s test were performed using IBM-SPSS statistical package version 22 (SPSS Inc., Chicago, IL, USA). A probability at *p* < 0.05 was considered statistically significant. For visualization of volatile metabolites and amino acid profiles, data were subjected to principal component analysis (PCA) in Multi-Experiment Viewer (MeV) version 4.8 (www.tm4.org/mev/) (13).

## Acknowledgements

The authors are grateful to the Research Program for Development of Small and Medium Enterprise, Researchers from the National Research Council of Thailand, Thailand Research Fund and Thailand Research Organizations Network (TRON) of years 2016 and 2019, for providing research fund, as well as Ratchadapisek Somphot Fund for the Postdoctoral Fellowship, and Chulalongkorn University in supporting Rachatida Det-udom^1^, Ph.D to work under this project.

## References

1. Phewpan A, Phuwaprisirisan P, Takahashi H, Ohshima C, Ngamchuachit P, Techaruvichit P, Dirndorfer S, Dawid C, Hofmann T, Keeratipibul S. 2020. Investigation of kokumi substances and bacteria in Thai fermented freshwater fish (Pla-ra). J Agric Food Chem 68:10345–10351. doi:10.1021/acs.jafc.9b06107

2. Ohshima C, Takahashi H, Insang S, Phraephaisarn C, Techaruvichit P, Khumthong R, Haraguchi H, Lopetcharat K, Keeratipibul S. 2019. Next-generation sequencing reveals predominant bacterial communities during fermentation of Thai fish sauce in large manufacturing plants. LWT 114:Article number 108375. doi:10.1016/j.lwt.2019.108375

3. Lee SH, Jung JY, Jeon CO. 2014. Microbial successions and metabolite changes during fermentation of salted shrimp (saeu-jeot) with different salt concentrations. Plos One 9:Article Number: e90115 doi:10.1371/journal.pone.0090115

4. Wang Z, Xu Z, Sun L, Dong L, Wang Z, Du M. 2020. Dynamics of microbial communities, texture and flavor in Suan zuo yu during fermentation. Food Chem 332:Article number 127364. doi:10.1016/j.foodchem.2020.127364

5. Guan L, Cho KH, Lee JH. 2011. Analysis of the cultivable bacterial community in jeotgal, a Korean salted and fermented seafood, and identification of its dominant bacteria. Food Microbiol 28:101–113. doi:10.1016/j.fm.2010.09.001

6. Jung MY, Kim TW, Lee C, Kim JY, Song HS, Kim YB, Ahn SW, Kim JS, Roh SW, Lee SH. 2018. Role of jeotgal, a Korean traditional fermented fish sauce, in microbial dynamics and metabolite profiles during kimchi fermentation. Food Chem 265:135–143. doi:10.1016/j.foodchem.2018.05.093

7. Md Zoqratt MZH, Gan HM. 2020. The microbiota of Malaysian fermented fish sauce. bioRxiv. doi:10.1101/2020.03.10.986513

8. Chhetri V, Prakitchaiwattana C, Settachaimongkon S. 2019. A potential protective culture; halophilic Bacillus isolates with bacteriocin encoding gene against Staphylococcus aureus in salt added foods. Food Control 104:292–299. doi:10.1016/j.foodcont.2019.04.043

9. Kim TW, Lee JH, Kim SE, Park MH, Chang HC, Kim HY. 2009. Analysis of microbial communities in doenjang, a Korean fermented soybean paste, using nested PCR-denaturing gradient gel electrophoresis. Int J Food Microbiol 131:265–271. doi:10.1016/j.ijfoodmicro.2009.03.001

10. Wei Q, Wang H, Chen Z, Lv Z, Xie Y, Lu F. 2013. Profiling of dynamic changes in the microbial community during the soy sauce fermentation process. Appl Microbiol Biotechnol 97:9111–9119. doi:10.1007/s00253-013-5146-9

11. Tanasupawat S, Thongsanit J, Okada S, Komagata K. 2002. Lactic acid bacteria isolated from soy sauce mash in Thailand. J Gen Appl Microbiol 48:201–209. doi:10.2323/jgam.48.201

12. Sánchez-Porro C, Martín S, Mellado E, Ventosa A. 2003. Diversity of moderately halophilic bacteria producing extracellular hydrolytic enzymes. J Appl Microbiol 94:295–300. doi:10.1046/j.1365-2672.2003.01834.x

13. Luangwilai M, Duangmal K, Chantaprasarn N, Settachaimongkon S. 2021. Comparative metabolite profiling of raw milk from subclinical and clinical mastitis cows using ^1^H-NMR combined with chemometric analysis. Int J Food Sci Tech 56:493–503. doi:10.1111/ijfs.14665

14. Tyn MT. 2004. Industrialization of Myanmar fish paste and sauce fermentation, p 737–762. *In* Steinkraus KH (ed), Industrialization of Indigenous Fermented Foods 2nd ed.

15. Prakitchaiwattana CJ, Fleet GH, Heard GM. 2004. Application and evaluation of denaturing gradient gel electrophoresis to analyse the yeast ecology of wine grapes. FEMS Yeast Res 4:865–877. doi:10.1016/j.femsyr.2004.05.004

16. Nakano MM, Dailly YP, Zuber P, Clark DP. 1997. Characterization of anaerobic fermentative growth of *Bacillus subtilis*: Identification of fermentation end products and genes required for growth. J Bacteriol 179:6749–6755. doi:10.1128/jb.179.21.6749-6755.1997

17. Kobayashi K, Shimojo S, Watanabe S. 2016. Contribution of a fermentation process using *Bacillus subtilis* (natto) to high polyamine contents of natto, a traditional Japanese fermented soy food. Food Sci Technol Res 22:153–157. doi:10.3136/fstr.22.153

18. Booker AE, Borton MA, Daly RA, Welch SA, Nicora CD, Hoyt DW, Wilson T, Purvine SO, Wolfe RA, Sharma S, Mouser PJ, Cole DR, Lipton MS, Wrighton KC, Wilkins MJ. 2017. Sulfide generation by dominant *Halanaerobium* microorganisms in hydraulically fractured shales. mSphere 2:Article number e00257-17. doi:10.1128/mSphere.00257-17

19. Booncharoen A, Visessanguan W, Kuncharoen N, Yiamsombut S, Santiyanont P, Mhuantong W, Charoensri S, Rojsitthisak P, Tanasupawat S. 2019. *Lentibacillus lipolyticus* sp. nov., a moderately halophilic bacterium isolated from shrimp paste (Ka-pi). Int J Syst Evol Microbiol 69:3529–3536. doi:10.1099/ijsem.0.003658

20. Pakdeeto A, Tanasupawat S, Thawai C, Moonmangmee S, Kudo T, Itoh T. 2007. *Lentibacillus kapialis* sp. nov., from fermented shrimp paste in Thailand. Int J Syst Evol Microbiol 57:364–369. doi:10.1099/ijs.0.64315-0

21. Satomi M, Kimura B, Mizoi M, Sato T, Fujii T. 1997. *Tetragenococcus muriaticus* sp. nov., a new moderately halophilic lactic acid bacterium isolated from fermented squid liver sauce. Int J Syst Bacteriol 47:832–836. doi:10.1099/00207713-47-3-832

22. Zhang L, Zhang L, Xu Y. 2020. Effects of *Tetragenococcus halophilus* and *Candida versatilis* on the production of aroma-active and umami-taste compounds during soy sauce fermentation. J Sci Food Agric 100:2782–2790. doi:10.1002/jsfa.10310

23. Zang J, Xu Y, Xia W, Regenstein JM. 2020. Quality, functionality, and microbiology of fermented fish: a review. Crit Rev Food Sci Nutr 60:1228–1242. doi:10.1080/10408398.2019.1565491

24. Xu Y, Zang J, Regenstein JM, Xia W. 2020. Technological roles of microorganisms in fish fermentation: a review. Crit Rev Food Sci Nutr:in press.

25. Kobayashi T, Kimura B, Fujii T. 2000. Strictly anaerobic halophiles isolated from canned Swedish fermented herrings (Surstromming). Int J Food Microbiol 54:81–89. doi:10.1016/S0168-1605(99)00172-5

26. Jung JY, Lee SH, Lee HJ, Jeon CO. 2013. Microbial succession and metabolite changes during fermentation of saeu-jeot: Traditional Korean salted seafood. Food Microbiol 34:360–368. doi:10.1016/j.fm.2013.01.009

27. Udomsil N, Chen S, Rodtong S, Yongsawatdigul J. 2017. Improvement of fish sauce quality by combined inoculation of *Tetragenococcus halophilus* MS33 and *Virgibacillus* sp. SK37. Food Control 73:930–938. doi:10.1016/j.foodcont.2016.10.007

28. Kaban G. 2013. Sucuk and pastirma: Microbiological changes and formation of volatile compounds. Meat Sci 95:912–918. doi:10.1016/j.meatsci.2013.03.021

29. Majumdar RK, Roy D, Bejjanki S, Bhaskar N. 2016. Chemical and microbial properties of shidal, a traditional fermented fish of Northeast India. J Food Sci Technol 53:401–410. doi:10.1007/s13197-015-1944-7

30. Lopetcharat K, Choi YJ, Park JW, Daeschel MA. 2001. Fish sauce products and manufacturing: A review. Food Rev Int 17:65–88. doi:10.1081/FRI-100000515

